## Supplementary Text File for "iPRESTO: automated discovery of biosynthetic sub-clusters linked to specific natural product substructures"

### Supplementary methods

#### Tokenising BGCs

To represent sequence similarity, BGCs were tokenised by converting them into strings of Pfam domains, using the HMMER3 tool hmmscan and the Pfam database version 32.0 [1, 2]. As Pfams are broad domain models, we divided the Pfams that are most important for BGCs into more specific domain models called ‘subPfams’, to increase the resolution for sub-cluster detection. To create subPfams, a Pfam is divided into more narrow domains models that cover the subspaces of that Pfam, by extracting the multiple sequence alignment of a Pfam and separating it into clades.  A new profile Hidden Markov Model (pHMM) is then built for each clade, each of which constitutes a subPfam (Fig 6A). The 112 biosynthetic Pfams that are most abundant in the antiSMASH database were converted into subPfams (S3 File). We subsequently used an pHMM database where these 112 Pfams were replaced with their corresponding subPfams in the Pfam database version 32.0. To query a BGC, we used hmmscan to scan against our pHMM database with the tc-cutoff as a cutoff on the bit score. Multiple hits in a gene were allowed to overlap by 10%. If the overlap was higher, only the hit with the highest bit score was kept. Following this approach, we tokenised each BGC as a string of genes, where each gene is a token represented as a combination of the present (sub)Pfam domains (S1B Fig).

#### Filtering redundant BGCs

To reduce phylogenetic bias, we filtered out redundant BGCs by constructing a similarity network of BGCs and choosing representative nodes from this network. As a similarity measure between BGCs, we used an Adjacency Index of domains (AI), which has been used previously to assess BGC similarity [3]. The AI between BGCs is calculated by dividing the number of all distinct shared pairs of adjacent domains by the total number of distinct pairs of adjacent domains, while ignoring gene boundaries. We constructed undirected graphs of similar BGCs by connecting two BGCs if their AI was above 0.95. We also connected two BGCs if one BGCs was fully contained in the other. To select representative BGCs from the graphs, all maximal cliques in the graph are identified using find_cliques from the networkx module, which is based on the algorithm described by Bron et al. [4]. Then, the BGC with the most domains is chosen from each maximal clique to remain in the analysis, iterating over the cliques from largest to smallest until there are no cliques left. If there is more than one BGC to choose from, the BGC with the fewest connections is retained in the analysis to preserve as much information as possible. BGCs in a clique that are not selected are filtered out (S9 Fig). This process is repeated until there are no connections left between BGCs.

#### Filtering domains

As we are interested in groups of genes that are directly responsible for the biosynthesis of chemical substructures, we chose to only detect sub-clusters of biosynthetic genes. To only select such genes, we discarded all Pfams that were not present in a list of 1,839 biosynthetic Pfams. We compiled this list by collecting all 3,010 EC-associated Pfams from ECDomainMiner using the lowest threshold [5]. We discarded domains from this list if they did not occur within a set of existing pre-calculated BGCs used in Kautsar et al. [6], which consists of the antiSMASH database and several fungal and plant BGCs. This list was filtered further by searching for keywords like transporter or DNA-binding. We then added 50 manually curated biosynthetic domains to the list that were not part of ECDomainMiner but occur frequently in BGCs within the antiSMASH-DB database, resulting in a list of 1,839 biosynthetic domains (S3 File). Additionally, Pfams were removed before sub-cluster detection if they occurred less than three times throughout the dataset. Subsequently, we removed all BGCs that contained fewer than two non-empty genes as result of Pfam filtering.

#### Clustering statistical sub-clusters

As the statistical method results in many sub-clusters, we clustered them into sub-cluster families (SCFs) and the SCFs into sub-cluster clans (SCCs). To do so, we used the K-means algorithm implemented in scikit learn with k-means++ seeding [7, 8]. We represented all sub-clusters as a presence/absence matrix of the tokenised genes on which we ran K-means with 1,000 iterations and 20 restarts. To construct SCFs, we assessed the K-means clustering of different numbers for k. We chose a clustering based on the lowest within cluster sum-of-squares (WCSS), while keeping the number of families to a minimum and trying to avoid the formation of one big ‘hairball’ cluster with unrelated sub-clusters. In order to cluster the SCFs into SCCs, we clustered the centroids from the SCF clustering and assessed the clustering of different numbers for k in the same way as for the SCFs. We deemed an SCF to be meaningful if it had three genes that were present in at least 60% of the sub-clusters in the SCF. Additionally, we removed redundant sub-clusters from each SCF. We deemed a sub-cluster redundant if it had the same occurrence as a bigger sub-cluster in which it was contained completely.

#### Benchmarking against SubClusterBlast

The 127 SubClusterBlast sub-clusters were extracted from https://bitbucket.org/antismash/antismash/src/master/antismash/generic_modules/subclusterblast/subclusters.txt [9]. From the 127 validated sub-clusters, 109 had matching accessions in the MIBiG database. To see how many known sub-clusters we could identify, we calculated the overlap between all known sub-clusters and the putative sub-clusters from one of the detection methods. We defined an overlap as the number of genes (domain combinations) from a known sub-cluster that are present in a putative sub-cluster, divided by the number of genes in the known sub-cluster. We considered a known sub-cluster to be detected if there was at least one putative sub-cluster matching the known sub-cluster with an overlap above 0.6.

#### Sub-cluster annotation

The annotation of sub-cluster motifs or sub-cluster clans (SCCs) with substructures is still a manual task with low throughput, which is why we annotated 45 sub-cluster motifs and some SCCs so far. In order to assign a substructure to a sub-cluster motif or SCC, we looked at the sub-cluster motifs and SCCs present in MIBiG BGCs, as their structures are validated. We considered an annotation appropriate for a sub-cluster motif or SCC if it is present in multiple MIBiG BGCs that share a similar substructure, while the genes in the sub-cluster comply with their proposed function in literature (S1 File). The latter is more valid for sub-cluster motifs and SCCs encompassing previously known sub-clusters as the genes from known sub-clusters are typically experimentally validated. To visualise and inspect identified sub-clusters, we implemented and modified a BGC visualisation script from Navarro-Muñoz et al. [3] to allow for sub-cluster visualisation.

#### Linking orphan NPs to BGCs through sub-cluster motifs

To find orphan NPs that contain a substructure encoded by one of the annotated sub-cluster motifs, we used the substructure search function on NPAtlas by drawing the substructure of interest [10]. We downloaded the tab delimited output containing information about the strain in which the matching NPs from NPAtlas where present. Together with the taxonomic metadata from the antiSMASH-DB dataset, we used link_np_to_bgc_subcl_motifs.py from the iPRESTO codebase to find BGCs in those strains that contain the sub-cluster motifs encoding the substructure of interest. By investigating the other parts of the identified BGCs, like predicted scaffolds, and searching in literature, we determined whether the identified BGC could putatively encode the biosynthesis of a given NP from NPAtlas. For constructing phylogenetic trees, CORASON was used with default settings on the sub-cluster regions [3]. For the identification of the akashin BGC, we ran antiSMASH 6.1.0 on the genome of *Streptomyces sp. F001* (QZWF00000000), and ran iPRESTO using the (annotated) sub-cluster motifs from our final model.

#### Correlation analysis in substructure-based integrative omics analysis

In order to correlate substructures to sub-clusters in a systematic manner, we used the Streptomyces/Salinispora dataset to link substructure models to the two different sub-cluster models derived in this research, using a previously defined correlation metric [11]. The substructure models constitute 300 Mass2Motifs discovered previously with the MS2LDA tool, based on MS/MS data from the Streptomyces/Salinispora dataset [12]. The two sub-cluster models were generated by querying all tokenised Streptomyces/Salinispora BGCs to the LDA model trained on the whole antiSMASH database dataset (PRESTO-TOP), and to the SCCs generated from the whole antiSMASH database dataset (PRESTO-STAT), respectively. A Boolean vector was created for each Mass2Motif, sub-cluster motif and SCC, representing the presence/absence patterns in all strains of the Streptomyces/Salinispora dataset. We excluded Mass2Motifs or clans if they were present in fewer than two strains. Each pair of Mass2Motif and sub-cluster motif or SCC was scored for a mutual presence/absence pattern across strains following the correlation metric proposed by Dorogazhi et al. [11] This correlation score constitutes scoring +10 if both members of a pair are present in a strain, +1 if both members of a pair are absent in a strain, -10 if the Mass2Motif is present in a strain while a sub-cluster motif or SCC is not, or 0 if the Mass2Motif is absent in a strain while a sub-cluster motif or SCC is present. We prioritised potentially valuable pairs by assessing how meaningful a positive score is in two ways: by calculating the maximum possible correlation score without changing the occurrences and by performing a permutation test. The permutation test was carried out by scrambling each Boolean vector 10,000 times, calculating 10,000 random scores for each pair and dividing the times a higher or equal score than the observed score occurs by 10,000.

### Supplementary tables

**Table A.** **Number of BGCs in the different datasets during different processing steps before sub-cluster detection.** The antiSMASH-DB dataset is a combination of the antiSMASH database, the MIBiG database v1.4 and the Streptomyces/Salinispora dataset, which is the dataset we used for the paper. The small dataset refers to the combination of the Streptomyces/Salinispora dataset and the MIBiG database, which we used for visualizing the graph-based redundancy filtering step.

| **Number of BGCs** | **antiSMASH-DB** | **MIBiG 1.4** | **Streptomyces/Salinispora** | **antiSMASH-DB dataset** | **Small dataset** |
| --- | --- | --- | --- | --- | --- |
| **Initial** | 152,122 | 1,819 | 5,927 | 159,868 | 7,746 |
| **On contig edge** | 41,914 | 0 | 1,367 | 43,281 | 1,367 |
| **Filtered** | 50,296 | 317 | 3,113 | 56,559 | 3,456 |
| **Final** | 59,912 | 1,502 | 1,447 | 60,028 | 2,923 |

**Table B.** **Equations used by PRESTO-STAT.**

| 1. Hypergeometric equation for adjacency interactions between gene A and gene B. B_1_: gene B not adjacent to gene A, B_2_: gene B adjacent to gene A, B_3_: gene B adjacent to gene A on both sides. N_1_, N_2_ and N_3_ represent all available positions in these three categories, while N_tot_ represent all positions, and B_tot_ all occurrences of gene B. | $\boldsymbol{P}_{\boldsymbol{d}}\boldsymbol{=}\frac{\left( \begin{matrix} \boldsymbol{N}_{\boldsymbol{1}} \\ \boldsymbol{B}_{\boldsymbol{1}} \end{matrix} \right)\left( \begin{matrix} \boldsymbol{N}_{\boldsymbol{2}} \\ \boldsymbol{B}_{\boldsymbol{2}} \end{matrix} \right)\left( \begin{matrix} \boldsymbol{N}_{\boldsymbol{3}} \\ \boldsymbol{B}_{\boldsymbol{3}} \end{matrix} \right)}{\left( \begin{matrix} \boldsymbol{N}_{\boldsymbol{tot}} \\ \boldsymbol{B}_{\boldsymbol{tot}} \end{matrix} \right)}$ |
| --- | --- |
| 2. Hypergeometric equation for co-localisation interactions between gene A and gene B. B_1_: gene B not co-localised with gene A, B_2_: gene B co-localised to gene A, B_nmax_: gene B co-localised with nmax gene A. N_1_, N_2_ and N_nmax_ represent all available positions in these three categories, while N_tot_ represent all positions, and B_tot_ all occurrences of gene B. | $P_{d}=\frac{\left( \begin{matrix} N_{1} \\ B_{1} \end{matrix} \right)\left( \begin{matrix} N_{2} \\ B_{2} \end{matrix} \right)\ldots\left( \begin{matrix} N_{n_{max}} \\ B_{n_{max}} \end{matrix} \right)}{\left( \begin{matrix} N_{tot} \\ B_{tot} \end{matrix} \right)}$ |
| 3. Simplified hypergeometric equation for co-localisation interactions between gene A and gene B. B_1_: gene B not co-localised with gene A, B_2_: gene B co-localised to gene A. N_1_ and N_2_ represent all available positions in these two categories, while N_tot_ represent all positions, and B_tot_ all occurrences of gene B. | $P_{d}=\frac{\left( \begin{matrix} N_{1} \\ B_{1} \end{matrix} \right)\left( \begin{matrix} N_{2} \\ B_{2} \end{matrix} \right)}{\left( \begin{matrix} N_{tot} \\ B_{tot} \end{matrix} \right)}$ |
| 4. Calculation of the p-value for an interaction. i: amount of interaction, i_obs_: observed amount of interaction. | $p=P_{i \geq i_{obs}}=1-P_{i \leq i_{obs}}=1-\sum_{i \leq i_{obs}} P_{d}$ |

### References supplementary information

7.	Arthur D, Vassilvitskii S, editors. k-means++: The advantages of careful seeding. Proceedings of the eighteenth annual ACM-SIAM symposium on Discrete algorithms; 2007: Society for Industrial and Applied Mathematics.
